# Bivalency of natalizumab promotes inhibition of dynamic VLA-4 adhesion beyond affinity gain

**DOI:** 10.1101/2025.07.16.665053

**Authors:** Marie-Pierre Valignat, Martine Biarnes, Dominique Touchard, Audrey Ricco, Patrick Chames, Olivier Theodoly, Philippe Robert

## Abstract

Natalizumab, a monoclonal IgG4 antibody used in the treatment of multiple sclerosis (MS), inhibits VLA-4 binding to VCAM-1, thereby reducing leukocyte recruitment to inflamed tissues. The intricate mechanisms underlying these effects remain unclear, particularly concerning the heavy-chain shuffling of IgG4. We conducted an in vitro study to quantify the impact of bivalent IgG and monovalent Fab forms of natalizumab on the capture and migration of human primary memory T lymphocytes under shear stress, using VCAM-1 and SDF-1α-coated surfaces. IgG natalizumab at concentrations near its cell surface EC50 showed significant of capture and resistance of adherent cells to shear stress, whereas significantly higher doses of Fab natalizumab, up to 100-fold greater than its cell surface EC50, were needed to achieve similar effects. These findings highlight that receptor occupancy alone may not adequately predict the functional outcomes of inhibitor antibodies. For optimal therapeutic effect, inhibition of cell-surface adhesion may require specific kinetic and geometric binding properties of antibodies.

## Introduction

Multiple sclerosis is a chronic inflammatory disease of the central nervous system where demyelinization followed by degeneration of axons results in nerve conduction blockade with physical, cognitive and sometimes psychiatric disabilities. Natalizumab, a recombinant humanized monoclonal antibody, targets the alpha-4 subunit of the α4β1 and α4β7 integrins present on leukocytes. It seeks to reduce central nervous system inflammation by decreasing leukocyte trafficking toward inflamed tissues(1–3). Natalizumab was the first therapeutic antibody approved for multiple sclerosis(4–8), and is also approved for treatment of Crohn’s disease. However, this treatment is not specific for central nervous system inflammation, it is prone to side effects due to the induced immunodepression (9–11), and its molecular mechanisms are not entirely understood(12). A better understanding of the mechanisms underlying the therapeutic effect of natalizumab could guide design of new therapeutic antibodies targeting adhesion molecules.

Natalizumab is a divalent IgG4 antibody that can undergo heavy-chain shuffling in plasma, which may transform its functional behavior into that of a monovalent antibody (13–16). Springer et al showed that the Fab variant of natalizumab functions as an allosteric inhibitor, operating not by obstructing the VCAM-1 binding site but by binding to an epitope in close proximity(17). VLA-4, like other integrins, exists in three distinct conformational states: the low-affinity bent-closed (BC) and extended-closed (EC) conformations, alongside the high-affinity extended-open conformation (EO)(18, 19). This ensemble undergoes an intricate equilibrium, finely tuned by allosteric regulations. Extracellular ligand binding, intracellular adaptor/inhibitor interactions, and tensile forces applied by the actin cytoskeleton on the integrin β-subunit collectively influence this equilibrium(20–23). The EO conformation emerges as the ultimate competent state for mediating cell adhesion, and Springer et al (17) used Mn2+ to induce the extended-open (EO) conformation representing the active form of the integrin with high ligand-binding affinity. They found that the Fab variant of natalizumab induced a notable reduction in the activity of Mn2+-activated VLA-4 for VCAM-1, shifting it from 1.9 µM to 23 µM, which was interpreted as a change in the conformational space accessible to the D2 domain of VCAM-1.

Various functional studies have investigated the ability of natalizumab to block the integrin Very Late Antigen-4 (VLA-4; α4β1; CD49d/CD29) expressed by leukocytes and reticulocytes in the context of sickle cell disease(24, 25). White et al.(24) used cytometry and flow assay experiments to demonstrate that adhesion to immobilized vascular cell adhesion molecule 1 (VCAM-1) versus natalizumab was dose dependent. They noticed a discrepancy between the natalizumab concentration for 50% occupation of VCAM-1 sites on cells (EC-50) and for 50% of cell resisting deadhesion under flow on VCAM-1 coated surfaces (IC-50), and concluded that a fraction of molecular occupancy is sufficient to induce cell deadhesion. They also compared the impact of natalizumab, which is bivalent, with its monomeric IgG4 counterpart. While the affinity of natalizumab to VLA-4 presented by reticulocytic cells was increased by a factor 6 as compared to monomeric IgG4, the inhibition of cell adhesion was increased by a factor 20. However, these in vitro study do not consider the pivotal role of surface-bound chemokines recently highlighted by Alon et al. and Sosa-Costa et al.(26) in modulating the EO conformation of VLA-4, and consequently cell adhesion and diapedesis.

In the present study, we probed in vitro the effect of natalizumab on T lymphocyte functions mediated by VLA-4 integrin in combination with the chemokine CXCL12. We also investigated the impact of shuffling by employing natalizumab Fab as a model for a shuffled and functionally monovalent antibody. We employed two biophysical methods to evaluate lymphocyte adhesion and migration on model substrates featuring VCAM-1 and CXCL12. We analysed the initial leukocyte adhesion step to activated endothelium under low shear conditions, and we replicated the migration step of leukocytes on activated endothelium under physiological shear flow. These two approaches allowed the detection of single integrin ligand bonds, and assessment of resistance to shear flow and cell migration velocity. Focusing on model experiment with key player VCAM-1 and CXCL12, our study contributes to new insights into the intricate interplay between chemokines, integrins, and the therapeutic potential of antibodies in regulating immune cell trafficking.

## Materials and methods

### Human T lymphocytes preparation

Human whole blood, collected in accordance with French law from healthy volunteers by Etablissement Francais du Sang, underwent gradient centrifugation to separate peripheral blood mononuclear cells. Primary memory T lymphocytes were negatively isolated using a Miltenyi Biotec pan-T cell sorting kit (reference 130-096-535, Miltenyi) with additional CD45RA MicroBeads (reference 130-045-901, Miltenyi). Subsequently, these cells were rested systematically overnight and maintained in complemented RPMI medium until further use.

Effector T lymphocytes were derived from primary memory T lymphocytes through incubation in complemented RPMI medium containing T Cell TransAct™ (Miltenyi Biotec, Bergish Gladbach, Germany), a polymeric nanomatrix conjugated to humanized CD3 and CD28 agonists. The cultivated T lymphocytes were then nurtured in RPMI 1640 (Gibco by Thermo Fischer Scientific, Waltham, MA), supplemented with 25 mM GlutaMax (Gibco by Thermo Fischer Scientific, Waltham, MA) and 10% fetal bovine serum (FBS; Gibco by Thermo Fischer Scientific, Waltham, MA) at 37°C, 5% CO2, in the presence of IL-2 (50 ng/ml; Miltenyi Biotec, Bergisch Gladbach, Germany). The cells were utilized 6 to 10 days post-stimulation.

### Molecules and surfaces preparation

Natalizumab was acquired from sterile vials previously utilized and sourced from Marseille University Hospitals, maintained in sterile conditions, and stored at 4°C. The preparation of Natalizumab Fab via papain cleavage was conducted using a dedicated kit (ThermoFisher Scientific, USA) following the manufacturer’s recommendations. Validation of the formation of enzymatic products with a uniform size was confirmed through western blotting, and the final concentration was determined using UV spectrophotometry. The chambers employed were uncoated Ibidi IV channel slides (Ibidi, Germany) measuring 12 mm in length and with a section of 0.1×1 mm².

For capture experiments, a solution of protein A (50 µg/ml, Sigma-Aldrich, France) and SDF-1α (1 µg/ml, Biotechne, USA) in phosphate-buffered saline (PBS) was deposited and allowed to incubate for 1 hour at 37°C. Following rinsing with PBS, a blocking solution of 4% bovine serum albumin (BSA, Sigma Aldrich, France) in PBS was added and incubated for 30 minutes at 37°C. After another PBS rinse, a solution of recombinant human VCAM-1-Fc chimera (10 µg/ml, Biotechne, USA) in PBS was added and left overnight at 4°C.

For adhesion experiments involving crawling T lymphocytes under flow conditions, a solution of VCAM-1-Fc (10 µg/ml, Biotechne, USA) and SDF-1α (1 µg/ml, Biotechne, USA) in PBS was deposited and incubated overnight at 4°C. Following a PBS rinse, a blocking solution of BSA in PBS was applied for 30 minutes at 37°C.

### Flow cytometry affinity assays

Primary memory T lymphocytes were incubated with various amounts of either IgG natalizumab or Fab natalizumab for 30 minutes in complemented RPMI medium. Free sites of VLA-4 integrins were then marked by a label-free anti-VLA-4 antibody, clone HP2/1 (Immunotech IM0764, Beckman Coulter, France). HP2/1 anti VLA-4 antibodies were revealed by incubation with a fluorescent anti-mouse secondary antibody (AF633 Life technologies reference A21052). Single cell fluorescence was measured using a LSR Fortessa X20 flow cytometer (BD Biosciences, USA); raw data were analysed using FlowJo software (BD Biosciences, USA).

### Cell-free binding assays of natalizumab to VLA-4

The quantification of Natalizumab-VLA-4 binding in a cell-free assay was conducted using a biolayer interferometry instrument, specifically the Octet R2 by Sartorius (BLI). Protein-G sensors were coated with a 10 µg/mL solution of IgG Natalizumab in PBS-BSA 0.1%. Solutions of Fc-fragment-free recombinant human VLA-4 heterodimer (obtained from R&D Systems, USA) were prepared in PBS-BSA 0.1%. The binding measurements were executed in accordance with the manufacturer’s recommended protocol.

### T Lymphocyte Capture Experiments

A specially prepared Ibidi chamber surface was connected to a Razel A-99 syringe pump (USA) and mounted on an inverted microscope (Olympus CK 40, Japan) equipped with a x10 objective, brightfield illumination, and a video camera (Sony, Japan) (Figure 1A). The chamber, pump syringe, and piping were filled with the designated amount of either IgG natalizumab or Fab natalizumab in RPMI medium, or RPMI medium alone for the negative control. The entire setup was enclosed in a custom thermoregulated box maintained at 37°C. After a 30-minute incubation with the chosen natalizumab type and concentration in RPMI medium, or RPMI medium alone for controls, approximately 2×10^5^ cells in 200 µl of medium were injected into the chamber.

**Figure 1.**
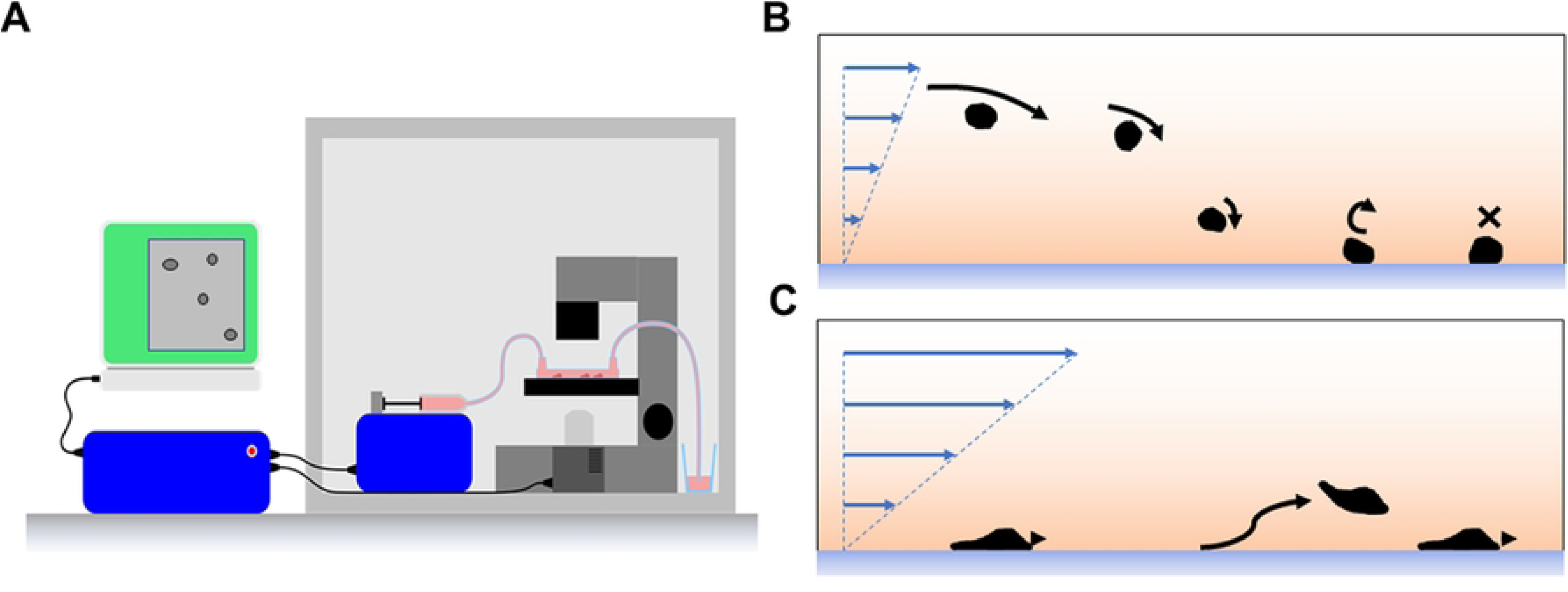
In panel **A**, in a thermoregulated chamber at 37°C (1), an inverted optical microscope (2) holds a laminar flow chamber connected to a syringe actuated by a syringe pump (3). A controller (4) adjust the syringe pump to obtain the desired shear stress in the flow chamber; a computer (5) acquires camera signal and retrieves cell trajectories, allowing to measure the number of cell arrest per distance travelled on the chamber surface and the duration of each arrest. For capture experiments (panel **B**), cells are injected in the chamber, sediment, roll and adhere to the chamber surface, under shear flows set from 0.003 Pa to 0.013 Pa. For adhesion of crawling T lymphocytes experiments (panel **C**), the cells that adhered during capture experiments shown in panel **B** migrated on the chamber surface and a physiological of 0.1 Pa shear stress was then applied to measure the velocity of cells and the fraction of cells withstanding a 10 minutes period of continuous shear stress.

The syringe pump applied a specified shear stress for five minutes while recording a movie at 25 frames per second. Shear stress was sequentially applied at 0.006 Pa, then at 0.003 Pa (Figure 1B). Between change in shear stress, a new suspension of cells was injected, effectively rinsing away any adhering cell from the previous condition.

### T Lymphocytes Adhesion of Crawling T Lymphocytes Experiments

The setup for T lymphocytes adhesion of crawling T lymphocytes experiments mirrored that of the lymphocyte capture experiment, with the only difference being the acquisition of movies at one frame every 10 seconds (Figure 1C). Adhesion assays for crawling T lymphocytes were conducted immediately following a 0.003 Pa capture assay. Cells adhered during the previous assay were subjected to a shear stress of 0.1 Pa for 10 minutes while being recorded.

## Data analysis

In T lymphocyte capture experiments, raw data were obtained for each shear flow and natalizumab concentration condition, presented as 25 frames per second M-JPEG compressed movies. A plug-in for ImageJ (NIH, USA), developed in Java (Oracle, USA), was written to extract the barycenter of each cell—detected through luminance thresholding—in every frame and subsequently reconstruct individual cell trajectories. Another plug-in was then utilized to generate velocity histograms for each cell, facilitating automated identification of cell arrests on the surface, measurement of arrest duration, and quantification of distances covered by cells sedimented on the surface. Linear binding density reflected association kinetics and was calculated as the ratio of the number of cell arrests to the total distance traveled by sedimented cells. Differences in dissociation kinetics were explored by pooling arrests durations for each experimental condition, then by building arrests survival curves representing the fraction of surviving arrests versus time.

In the case of cell adhesion experiments involving crawling T lymphocytes, raw data were collected for each shear flow and natalizumab concentration condition in the form of M-JPEG compressed movies. For each flow sequence, the number of cells in the initial image was tallied, and the motion of these cells was tracked across the subsequent 30 images. The percentage of migrating cells was then computed as the ratio of the remaining and migrating cells to the initial number of cells at the onset of the flow sequence. Velocities of individual migrating cells were computed and compared for the different experimental conditions.

## Results

### Determination of natalizumab binding properties in IgG and Fab forms

The characterization of IgG natalizumab binding to free VLA-4 was executed using a biolayer interferometry (BLI) setup, where sensor-bound IgG natalizumab and recombinant VLA-4 heterodimer in solution were employed (**Error! Reference source not found.**). This resulted in a monovalent binding affinity of 13 nM, aligning well with previously published data (27). To quantify the free VLA-4 sites in memory T lymphocytes or effector T lymphocytes under varying doses of IgG natalizumab and Fab natalizumab, the binding of fluorescent anti-VLA-4 antibody (clone HP2/1) was measured. Cells underwent incubation with IgG natalizumab doses ranging from 0.02 nM to 20 nM, or with Fab natalizumab doses spanning from 0.06 nM to 600 nM. Flow cytometry experiments (Figure 2) revealed half-occupancy of VLA-4 at a dosage of 0.4 nM with IgG natalizumab for both memory T lymphocytes and effector T lymphocytes. Similarly, half-occupancy was achieved at 11 nM and 14 nM with Fab natalizumab for memory T lymphocytes and effector T lymphocytes, respectively. In summary, on two different cell models, affinity of IgG natalizumab is approximatively 30 times higher than affinity of Fab natalizumab. These concentrations align with published data obtained with different cell types (17, 24) and represent the IgG or Fab binding affinity EC_50_ for cell-bound VLA-4.

**Figure 2.**
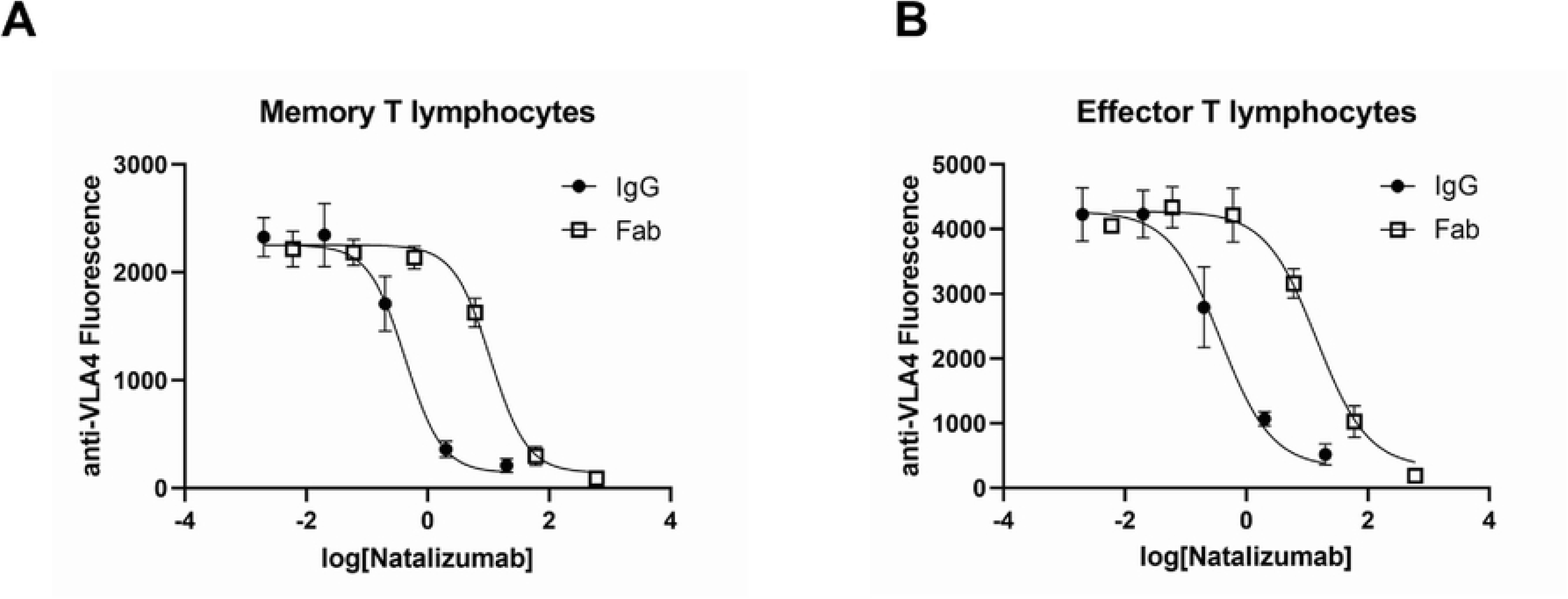
Flow cytometry cell-surface EC*_50_* measurements of IgG natalizumab and Fab natalizumab were obtained by quantifying the binding of the fluorescent anti-VLA-4 antibody (clone HP2/1) after incubating each type of cell with increasing amounts of IgG natalizumab or Fab natalizumab (fluorescence signal on the left axis, concentration on the bottom axis). In panel ***A***, measurements were performed on primary memory T lymphocytes, and in panel ***B***, measurements were performed on effector T lymphocytes.

### IgG natalizumab inhibits T lymphocytes capture and reduces cell arrest durations under flow on VCAM-1 at concentration close to its cell-surface *EC_50_* with more than 10-fold higher efficiency than Fab natalizumab

To assess *in vitro* the effect of natalizumab on initial capture of flowing cells by the endothelium, we quantified the arrest rate of sedimented cells as they flowed downstream by dividing the number of arrests by the distance they covered along the floor of a flow chamber before being captured. These capture experiments were performed under two different shear stresses (0.003 Pa and 0.006 Pa), allowing the probing of integrin-ligand interactions in the absence of selectin-PSGL-1 and other integrin-ligand interactions. Flow chamber surfaces were coated with saturating amounts of VCAM-1-Fc chimera on protein A and SDF-1α. Memory primary human T cells were incubated with IgG natalizumab at doses ranging from 0.02 nM to 20 nM, as well as Fab natalizumab at doses ranging from 0.06 nM to 600 nM prior to the experiments. IgG natalizumab and Fab natalizumab were also present in the flow medium during capture experiments, with corresponding amounts for each condition. Capture experiments with IgG natalizumab (Figure 3A) showed a significant decrease in adhesion of memory T lymphocytes, consistent with a decrease of VLA-4 affinity. This effect is statistically significant at doses as low as 0.02 nM under 0.003 Pa, corresponding to approximately a tenth of the IgG natalizumab binding affinity EC_50_. Additionally, the effect is observed at 0.2 nM under 0.006 Pa, aligning with the IgG natalizumab binding affinity EC50. On opposite, capture experiments with Fab natalizumab show significant inhibition at concentration of 60 nM or higher. This value is ten times greater than its binding affinity EC50 and is observed under both 0.003 Pa and 0.006 Pa shear stresses (Figure 3B). Altogether, effects on cell capture are significant for association kinetics with IgG natalizumab for amounts of antibody similar to its EC_50_while Fab natalizumab shows significant effect only for doses 10 to 100 times above cell surface EC_50_.

**Figure 3.**
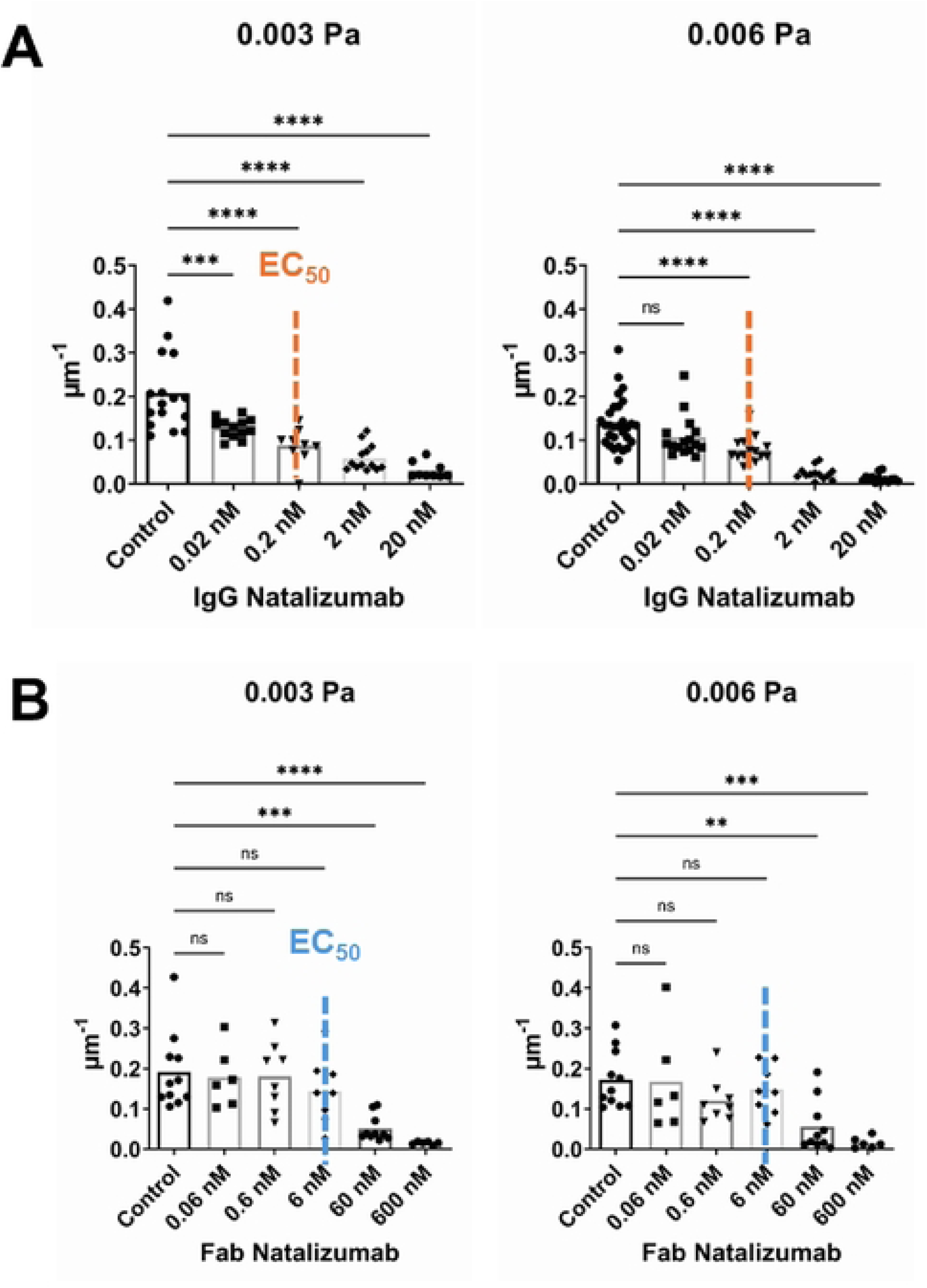
Capture experiments were performed by perfusing a suspension of primary human memory T lymphocytes in a flow chamber with its surface coated with VCAM-1 and SDF-1α. Values represent the number of cellular arrest events on the surface per the total distance travelled by sedimented cells, with each dot corresponding to an independent experiment. In panel **A**, cells were exposed to three different shear stresses (from left to right, 0.003 Pa, 0.006 Pa and 0.013 Pa) with concentrations of IgG natalizumab ranging from 0.02 nM to 20 nM (used to pre-incubate the cells and also present in the flow chamber medium). The orange dotted lines indicate the IgG natalizumab cell-surface EC_50_ value. In panel **B**, cells were exposed to three different shear stresses (from left to right, 0.003 Pa, 0.006 Pa and 0.013 Pa) with Fab natalizumab concentrations ranging from 0.06 nM to 60 nM (used to pre-incubate the cells and also present in the flow chamber medium). The blue dotted lines represent the Fab natalizumab cell-surface EC_50_ value. A dose-response effect of IgG natalizumab was observed at concentrations 10 times below the cell surface EC_50_ for all tested shear stresses. Fab natalizumab exhibited a significant effect only at doses 10 times above the cell surface EC_50_ for the two lower shear stresses. One-way ANOVA, followed by the Tukey multiple-comparison test, confirmed or infirmed the presence of significant differences for different values of Natalizumab concentrations (p values * <0.1, **<0.01, ***<0.001, ****<0.0001).

### IgG natalizumab inhibits adhesion of crawling T lymphocytes under physiological flow on VCAM-1 at concentration close to *EC_50_* with 100-fold higher efficiency than Fab natalizumab

The objective of our adhesion assays involving crawling T lymphocytes is to quantify cell adhesion to VCAM-1 on the chamber surface in presence of adsorbed CXCL12. This quantification is achieved by enumerating cells migrating before and after exposure to physiological shear flow. The experiments involved the tracking of memory primary human T cells adhered to the chamber surface following a 5-minute perfusion at 0.003 Pa. Preceding the experiment, cells were incubated with IgG natalizumab (at doses ranging from 0.02 nM to 200 nM) or Fab natalizumab (at doses ranging from 0.06 nM to 600 nM) for 30 minutes. Additionally, both IgG natalizumab and Fab natalizumab were added to the flow medium in corresponding amounts for each experiment. A shear stress of 0.1 Pa was applied, and cells were tracked for 10-minutes intervals for each treatment (Figure 4A). A significant and robust impact on the fraction of adherent cells was evident for IgG natalizumab, with effects observed at concentrations as low as 0.2 nM, closely aligning with its cell surface EC_50_. In contrast, Fab natalizumab exhibited its initial significant effect at 60 nM, and an effect comparable to IgG natalizumab at 0.2 nM was achieved at 600 nM, a concentration one hundred times higher than its cell surface EC_50_. A small increase in velocity was observed for higher amounts of IgG natalizumab and Fab natalizumab (Figure 4B).

**Figure 4.**
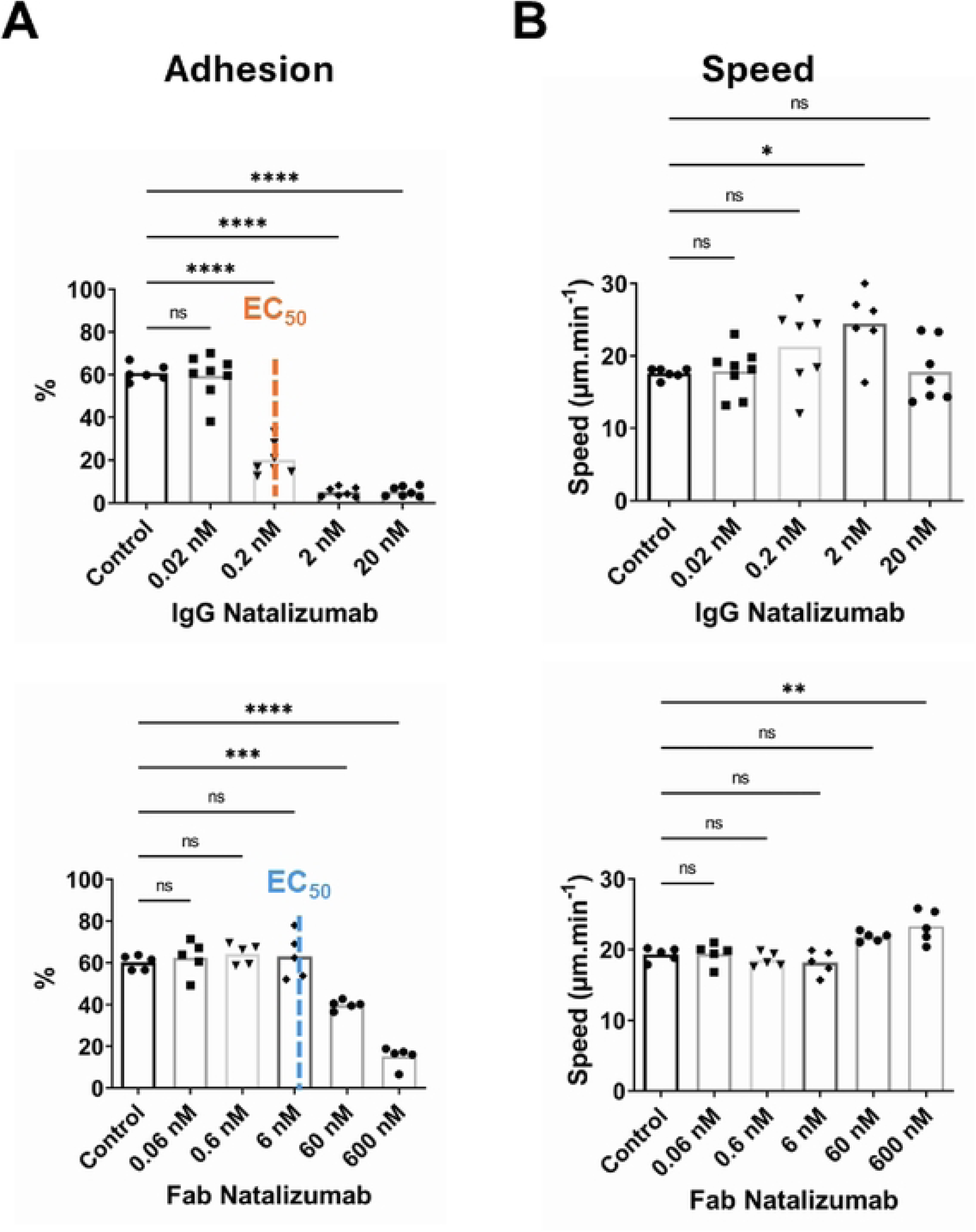
Adhesion of crawling T lymphocytes assays were performed by injecting a suspension of primary human memory T lymphocytes into a flow chamber with its surface coated with VCAM-1 and SDF-1α. In In panel **A**, values represent the proportion of cells remaining adhered after a 10-minutes period under 0.1 Pa shear stress, with each dot corresponding to an independent experiment. Left, cells were pre-incubated with concentrations of IgG natalizumab ranging from 0.02 nM to 20 nM, also present in the flow chamber medium during the experiment. An orange dotted line indicates the IgG natalizumab cell-surface EC_50_ value. Right, cells were pre-incubated with Fab natalizumab concentrations ranging from 0.06 nM to 60 nM, also present in the flow chamber medium during the experiment. A blue dotted line represents the Fab natalizumab cell-surface EC_50_ value. A 3-fold reduction in the proportion of adhering cells is observed for IgG natalizumab, occurring at concentrations around and above the EC_50_. In contrast, Fab natalizumab exhibits only a minor effect under similar shear stresses, noticeable only at doses approximately 10 times the EC_50_. To achieve a similar 3-fold reduction in the proportion of remaining adhered cells, a higher dose in the range of 100 times the EC_50_ is required. In panel **B**, values represent the average velocity of cells migrating under 0.1 Pa shear stress. Left, cells were pre-incubated with concentrations of IgG natalizumab ranging from 0.02 nM to 20 nM, also present in the flow chamber medium during the experiment. An orange dotted line indicates the IgG natalizumab cell-surface EC_50_ value. Right, cells were pre-incubated with Fab natalizumab concentrations ranging from 0.06 nM to 60 nM, also present in the flow chamber medium during the experiment. A blue dotted line represents the Fab natalizumab cell-surface EC_50_ value. A small effect of either IgG natalizumab or Fab natalizumab may be seen at higher amounts of the antibody.

## Discussion

We employed two biophysical methods measuring three different parameters to assess cell capture and adhesion of crawling cells. Capture experiments were able to detect the effect of natalizumab on very small numbers of integrin-ligand bonds, including single bonds. If a lymphocyte would bind to the surface through a single VLA-4/VCAM-1 interaction, a force of 2.5 and 5 pN would be applied on this bond for shear forces of 0.003 Pa and 0.006 Pa respectively, which is an easily observable range(28). On opposite, migration assays would quantify cell detachment under physiological shear forces and therefore the effect of natalizumab on cell-wide populations of integrins.

Our cell-surface affinity measurements of both forms of natalizumab are consistent with previous works: our results for Fab binding to T lymphocytes are in very good accordance with two previous work measuring binding of natalizumab Fab on lymphocytes(17) or reticulocyte surfaces, *i.e*, 6nM(24). Our IgG natalizumab affinity measurement at lymphocyte surface yields a value of 0.2nM also in good accordance with the 0.28nM value measured on lymphocytes(17), but lower than the value of 0.93nM observed on reticulocytes(24). This consistency of natalizumab affinity is remarkable knowing that some measurements are performed with Mn2+ to force integrins in high-affinity extended-open (EO) state(17), whereas we performed affinity measurement on memory T lymphocytes in suspension, for which most integrins are in the low-affinity bent-closed (BC) state. This suggests strongly that the affinity of Natalizumab is very similar for the integrins in open extended or bended closed states.

Our investigations also revealed a potent inhibitory effect of IgG natalizumab when employing antibody concentrations proximal to its EC_50_ measured on primary T lymphocytes, whereas Fab natalizumab necessitated concentrations 10 to 100 times higher than its EC_50_ to achieve a comparable inhibitory effect. If the affinity for VLA-4 were the sole determinant of natalizumab’s impact, IgG natalizumab and Fab natalizumab would exhibit identical effects at their respective EC_50_. Our findings therefore indicate that receptor occupancy alone does not elucidate the observed differences on VLA-4 mediated cell adhesion between the two forms of natalizumab.

A first mechanistic hypothesis is that IgG natalizumab might stabilize the bent or unbent state of two neighbouring VLA-4 molecules by solidarizing their extracellular domains. The unfolding of a bent VLA-4 could be hampered if its distal domain would be linked to the distal domain of a neighbouring bent VLA-4. Memory T lymphocytes, which predominantly maintain their integrins in a bent closed state while in suspension, would exhibit significantly reduced adhesion to VCAM-1/SDF-α substrates when treated with IgG natalizumab at EC50. This reduction in adhesion would occur because the VLA-4 molecules bound to IgG natalizumab would be locked in a non-affine state. In contrast, when treated with Fab natalizumab at EC50, the VLA-4 molecules could transition into an open conformation, allowing for greater adhesion (Figure 5A).

**Figure 5.**
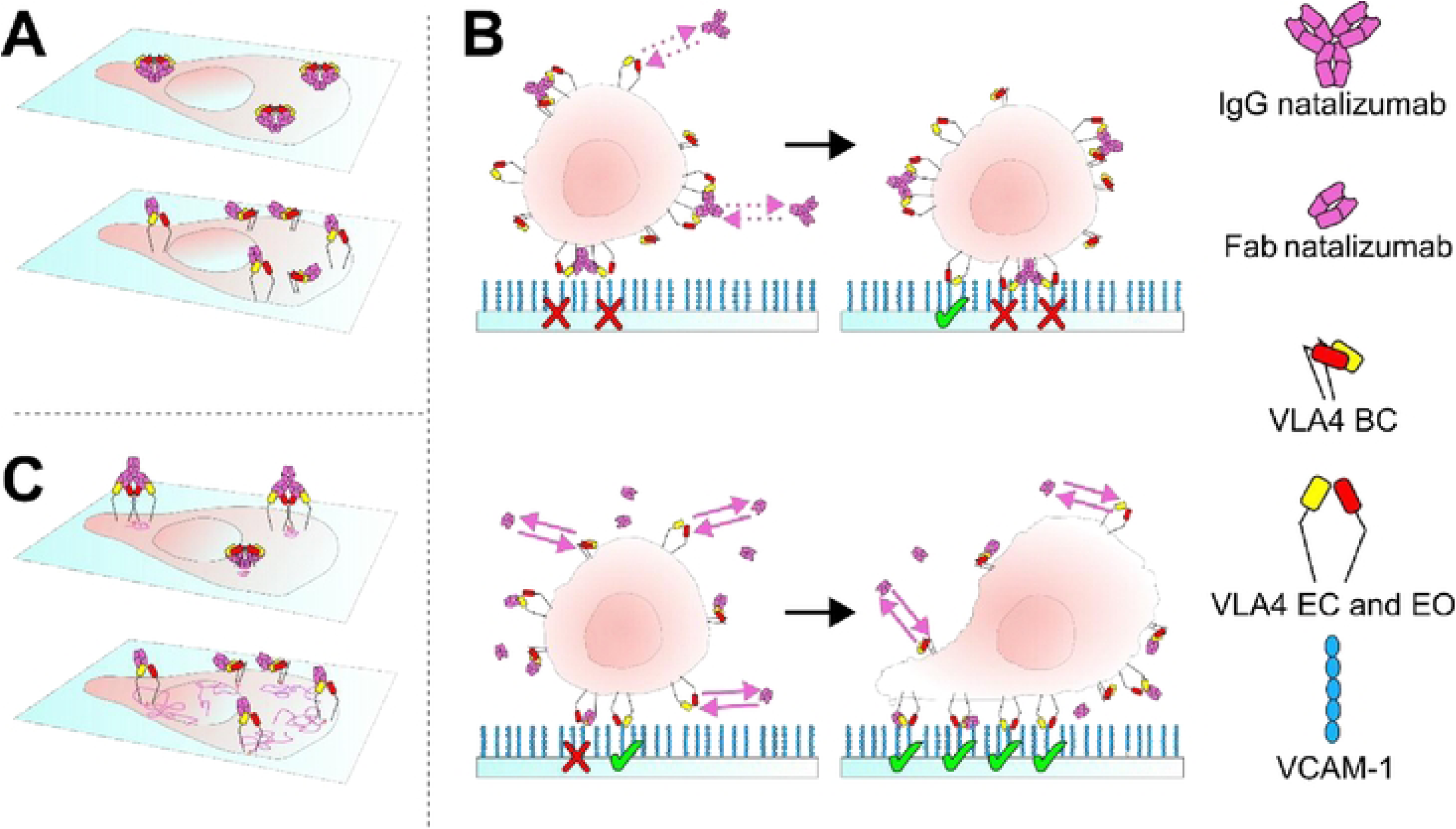
In panel **A**, we propose a first hypothetical effect of valency of natalizumab by acting on conformation of VLA-4 molecules. Top, IgG natalizumab is bound to two VLA-4 β_1_ chain in folded conformation, stabilizing both integrin in the folded state by hampering integrin head motions. Bottom, Fab natalizumab is bound to VLA-4 β_1_ chain with supposedly little effect on VLA-4 ability to undergo conformational change. In panel **B,** we propose a second hypothetical effect of valency of natalizumab. Top, IgG natalizumab binds with low dissociation kinetics to VLA-4, preventing adhesion of occupied VLA-4. Bottom, Fab natalizumab detaches faster from VLA-4, freeing VLA-4 for adhesion. If subsequent rebinding of natalizumab to VLA-4 does not promote deadhesion to VCAM-1, VLA-4 molecules may ultimately all adhere, effectively acting like a ratchet. In panel **C**, we propose a third hypothetical effect of valency of natalizumab by acting on membrane diffusion of VLA-4 molecules on a cell surface. Top, IgG natalizumab is bound to two VLA-4 β_1_ chain with supposedly reduction of diffusion area and velocity, reducing the efficiency of VLA-4 binding. Bottom, Fab natalizumab is bound to VLA-4 β_1_ chain with supposedly little effect on VLA-4 diffusion (represented by pink paths).

Alternatively, bivalency may play a pivotal role in cell adhesion by enhancing the sustained binding of IgG natalizumab to cell surface compared to Fab natalizumab. Due to local effects, the dissociation of a divalent IgG natalizumab from cell surface is expected to be slower than the dissociation of monovalent Fab natalizumab from cell surface(29). Besides, crystallography data suggested that natalizumab, either IgG or Fab, induced a reduction in VLA-4-VCAM-1 association kinetics rather than an increase in VLA-4-VCAM-1 dissociation kinetics(17). At the EC_50_ concentration, prior to the initial interaction of a cell with a surface, half of the VLA-4 receptors are not bound to natalizumab and can engage a surface coated by VCAM-1 with high intrinsic association kinetics, whereas the other half is bound to natalizumab and can engage a surface coated by VCAM-1 with reduced kinetics. In a static or instantaneous view, one therefore expects no major difference for the initial binding of a cell to a surface coated by VCAM-1 in presence of IgG or Fab natalizumab at EC50. However, since the half of VLA-4 receptors linked to natalizumab have a faster rate of exchange in the case of Fab than IgG, one expects over time that more VLA-4 will become unlinked from natalizumab in the Fab case, which will promote the creation of more additional bounds between the cell and a VCAM-1 coated surface. Hence, in the case of Fab natalizumab in solution, VLA-4 molecules would increasingly bind VCAM-1 in a ratchet effect, whereas in the case of IgG natalizumab presence in solution, the slower detachment of IgG natalizumab from VLA-4 molecules induces a lesser ratchet effect (Figure 5B). The difference of this dynamic ratchet effect is expected to be stronger for smaller duration of cell-surface contact, which is consistent with our observation that Fab natalizumab shows much weaker inhibition of capture events.

Dimerization of membrane molecules is also known to be critical in immunology to trigger signaling cascades(30). One might hypothesize that the binding of IgG natalizumab to VLA-4 triggers cell signalling distinct from that induced by the binding of Fab natalizumab. This difference could be attributed to the dimerization of VLA-4, leading to reduced effective adhesion. However, ligation of VLA-4 by antibodies has been described mostly as an activating signal in T lymphocytes (31, 32), in B lymphocytes(33), in eosinophils (34), and in monocytes-like THP1 cells(35, 36). Activated leukocytes usually display enhanced adhesion: anti-VLA-4 antibodies were reported to stimulate adhesion of T lymphocytes toward LFA-1 (37) or TCR (38). It appears therefore improbable that a signal resulting from antibody binding to VLA-4 would contribute to therapeutic effect by inhibition of adhesion.

Finally, VLA-4 molecules rapidly form clusters upon detection of SDF-1α by T lymphocytes(22). The regulation of their diffusion and clustering plays a critical role in cell adhesion and divalent binding may hinder this aspect of cell adhesion process by impeding the membrane diffusion of VLA-4 (Figure 5C).

The therapeutic administration of natalizumab is associated with plasma concentrations ranging from 18 µg/mL (120 nM) to 80 µg/mL (530 nM), depending on the dosing time, for a monthly 300 mg dose (39). In contrast, the IgG4 norm in patients typically hovers around 1 mg/mL (6 µM). A numerical model depicting IgG4 shuffling following the injection of a 300 mg dose of the therapeutic antibody indicates that bivalent IgG natalizumab levels in serum undergo a dynamic shift from 1 µg/mL (6 nM) to 0.1 µg/mL (0.6 nM) over the month following infusion(40). Concurrently, heterodimeric heavy-chain shuffled natalizumab/random IgG4 concentrations evolve from 70-80 µg/mL (500 nM) to 10 µg/mL (60 nM) during the same timeframe. Both concentrations prove to be substantially effective in functionally inhibiting VLA-4-VCAM-1 binding in plasma, as determined in our experimental model. According to pharmacokinetic data, the function of natalizumab in the central nervous system should be low since total natalizumab (IgG and Fab) concentrations in cerebrospinal fluid (CSF) are a hundred-fold to one thousand-fold below plasmatic levels (44ng/ml or 290 pM) (39). Our functional findings suggest that neither form of natalizumab achieves efficacy within the CSF.

IgG4 exhibits minimal or negligible interactions with Fc receptors and molecules related to humoral immunity, rendering them promising candidates for therapeutic antibodies with blocking capabilities. However, heavy-chain shuffling can transform them into functional monomers, leading to a decrease in the overall immunoglobulin’s affinity unless modifications are implemented by drug designers to mitigate this effect. Bivalency may benefit a therapeutic antibody by increasing its affinity toward a target. Our research suggests that therapeutic antibodies targeting adhesion molecules involved in cell-cell interactions could benefit further from bivalency. This would be attributed to the specific kinetics of cell-cell adhesion. Cell-cell or cell-surface adhesion is an obvious and important step of leukocyte recruitment, but also of many critical features of the immune response, both for signaling and effector function: antigen presentation to T lymphocytes, B lymphocyte activation by T lymphocytes, ADCC, ADPC occur through cell-cell or cell-pathogen surface interactions. All are key functions of the immune response, involved in pathology, making them important targets for therapeutic control through therapeutic antibodies(41).

Antibodies currently designed with anti-shuffling modifications could already be capitalizing on the advantages of bivalency(42). In the context of a modified, functionally bivalent natalizumab, lower injected doses might prove effective, and even the limited quantity present in cerebrospinal fluid could potentially yield added clinical benefits.

## Acknowledgments

1. D. Touchard passed away before the submission of the final version of this manuscript. M.-P. Valignat accepts responsibility for the integrity and validity of the data collected and analyzed. This work was supported by Agence Nationale de la Recherche (grants RECRUTE AAP CE15); LABEX INFORM; Région PACA and the Turing Centre for Living systems. Also, it has been carried out with the financial support of the Regional Council of Provence-Alpes-Côte d’Azur and with the financial support of the A*MIDEX (n° ANR-11-IDEX-0001-02), funded by the Investissements d’Avenir project funded by the French Government, managed by the French National Research Agency (ANR).

## References

1. G. P. A. Rice, H.-P. Hartung, P. A. Calabresi, Anti-alpha4 integrin therapy for multiple sclerosis: mechanisms and rationale. Neurology 64, 1336–1342 (2005).

2. J. L. Baron, J. A. Madri, N. H. Ruddle, G. Hashim, C. A. Janeway, Surface expression of alpha 4 integrin by CD4 T cells is required for their entry into brain parenchyma. J Exp Med 177, 57–68 (1993).

3. A. Prat, et al., Migration of multiple sclerosis lymphocytes through brain endothelium. Archives of neurology 59, 391–397 (2002).

4. D. H. Miller, et al., A controlled trial of natalizumab for relapsing multiple sclerosis. N Engl J Med 348, 15–23 (2003).

5. C. H. Polman, et al., A Randomized, Placebo-Controlled Trial of Natalizumab for Relapsing Multiple Sclerosis. New England Journal of Medicine 354, 899–910 (2006).

6. R. A. Rudick, et al., Natalizumab plus interferon beta-1a for relapsing multiple sclerosis. N Engl J Med 354, 911–923 (2006).

7. P. L. McCormack, Natalizumab: a review of its use in the management of relapsing-remitting multiple sclerosis. Drugs 73, 1463–1481 (2013).

8. P. O’Connor, et al., Relapse rates and enhancing lesions in a phase II trial of natalizumab in multiple sclerosis. Mult Scler 11, 568–572 (2005).

9. G. P. A. Rice, H.-P. Hartung, P. A. Calabresi, Anti-alpha4 integrin therapy for multiple sclerosis: mechanisms and rationale. Neurology 64, 1336–1342 (2005).

10. P. L. McCormack, Natalizumab: a review of its use in the management of relapsing-remitting multiple sclerosis. Drugs 73, 1463–1481 (2013).

11. M. Tintore, A. Vidal-Jordana, J. Sastre-Garriga, Treatment of multiple sclerosis - success from bench to bedside. Nat Rev Neurol 15, 53–58 (2019).

12. C. Coisne, W. Mao, B. Engelhardt, Cutting edge: Natalizumab blocks adhesion but not initial contact of human T cells to the blood-brain barrier in vivo in an animal model of multiple sclerosis. J Immunol 182, 5909–5913 (2009).

13. F. Debaene, et al., Time resolved native ion-mobility mass spectrometry to monitor dynamics of IgG4 Fab arm exchange and “bispecific” monoclonal antibody formation. Anal Chem 85, 9785–9792 (2013).

14. M. van der Neut Kolfschoten, et al., Anti-inflammatory activity of human IgG4 antibodies by dynamic Fab arm exchange. Science 317, 1554–1557 (2007).

15. J. Schuurman, et al., Normal human immunoglobulin G4 is bispecific: it has two different antigen-combining sites. Immunology 97, 693–698 (1999).

16. A. F. Labrijn, et al., Therapeutic IgG4 antibodies engage in Fab-arm exchange with endogenous human IgG4 in vivo. Nat Biotechnol 27, 767–771 (2009).

17. Y. Yu, T. Schürpf, T. A. Springer, How Natalizumab Binds and Antagonizes α4 Integrins*. Journal of Biological Chemistry 288, 32314–32325 (2013).

18. A. Chigaev, L. A. Sklar, Aspects of VLA-4 and LFA-1 regulation that may contribute to rolling and firm adhesion. Front Immunol 3, 242 (2012).

19. A. Chigaev, et al., Alpha4beta1 integrin affinity changes govern cell adhesion. J Biol Chem 278, 38174–38182 (2003).

20. A. E. May, F. J. Neumann, A. Schömig, K. T. Preissner, VLA-4 (alpha(4)beta(1)) engagement defines a novel activation pathway for beta(2) integrin-dependent leukocyte adhesion involving the urokinase receptor. Blood 96, 506–513 (2000).

21. Y. Nojima, D. M. Rothstein, K. Sugita, S. F. Schlossman, C. Morimoto, Ligation of VLA-4 on T cells stimulates tyrosine phosphorylation of a 105-kD protein. J Exp Med 175, 1045–1053 (1992).

22. V. Grabovsky, et al., Subsecond induction of alpha4 integrin clustering by immobilized chemokines stimulates leukocyte tethering and rolling on endothelial vascular cell adhesion molecule 1 under flow conditions. J Exp Med 192, 495–506 (2000).

23. Functional Mapping of Adhesiveness on Live Cells Reveals How Guidance Phenotypes Can Emerge From Complex Spatiotemporal Integrin Regulation - PubMed. Available at: https://pubmed-ncbi-nlm-nih-gov.proxy.insermbiblio.inist.fr/33898401/ [Accessed 6 August 2024].

24. J. White, et al., VLA-4 blockade by natalizumab inhibits sickle reticulocyte and leucocyte adhesion during simulated blood flow. British Journal of Haematology 174, 970–982 (2016).

25. P. C. Hines, et al., Natalizumab Blocks VLA-4 Mediated Red Blood Cell Adhesion and Is a Potential Therapy for Sickle Cell Disease. Blood 124, 221 (2014).

26. A. Sosa-Costa, et al., Lateral Mobility and Nanoscale Spatial Arrangement of Chemokine-activated α4β1 Integrins on T Cells. Journal of Biological Chemistry 291, 21053–21062 (2016).

28. X. Zhang, S. E. Craig, H. Kirby, M. J. Humphries, V. T. Moy, Molecular basis for the dynamic strength of the integrin alpha4beta1/VCAM-1 interaction. Biophys J 87, 3470–3478 (2004).

29. E. N. Kaufman, R. K. Jain, Effect of Bivalent Interaction upon Apparent Antibody Affinity: Experimental Confirmation of Theory Using Fluorescence Photobleaching and Implications for Antibody Binding Assays.

30. M. Mellado, J. M. Rodríguez-Frade, S. Mañes, C. Martínez-A, Chemokine Signaling and Functional Responses: The Role of Receptor Dimerization and TK Pathway Activation. Annual Review of Immunology 19, 397–421 (2001).

31. Y. Nojima, D. M. Rothstein, K. Sugita, S. F. Schlossman, C. Morimoto, Ligation of VLA-4 on T cells stimulates tyrosine phosphorylation of a 105-kD protein. J Exp Med 175, 1045–1053 (1992).

32. I. Ricard, M. D. Payet, G. Dupuis, Clustering the adhesion molecules VLA-4 (CD49d/CD29) in Jurkat T cells or VCAM-1 (CD106) in endothelial (ECV 304) cells activates the phosphoinositide pathway and triggers Ca2+ mobilization. Eur J Immunol 27, 1530–1538 (1997).

33. S. N. Manie, et al., Stimulation of tyrosine phosphorylation after ligation of beta7 and beta1 integrins on human B cells. Blood 87, 1855–1861 (1996).

35. M. Colden-Stanfield, Clustering of very late antigen-4 integrins modulates K(+) currents to alter Ca(2+)-mediated monocyte function. Am J Physiol Cell Physiol 283, C990–C1000 (2002).

36. I. D. McGilvray, et al., VLA-4 integrin cross-linking on human monocytic THP-1 cells induces tissue factor expression by a mechanism involving mitogen-activated protein kinase. J Biol Chem 272, 10287–10294 (1997).

37. A. E. May, F. J. Neumann, A. Schömig, K. T. Preissner, VLA-4 (alpha(4)beta(1)) engagement defines a novel activation pathway for beta(2) integrin-dependent leukocyte adhesion involving the urokinase receptor. Blood 96, 506–513 (2000).

38. A.-M. Cimo, Z. Ahmed, B. W. McIntyre, D. E. Lewis, J. E. Ladbury, CD25 and CD69 induction by α4β1 outside-in signalling requires TCR early signalling complex proteins. The Biochemical journal 454, 109–121 (2013).

39. T. Sehr, et al., New insights into the pharmacokinetics and pharmacodynamics of natalizumab treatment for patients with multiple sclerosis, obtained from clinical and in vitro studies. J Neuroinflammation 13, 164 (2016).

40. T. Sehr, et al., New insights into the pharmacokinetics and pharmacodynamics of natalizumab treatment for patients with multiple sclerosis, obtained from clinical and in vitro studies. J Neuroinflammation 13, 164 (2016).

41. R. J. Slack, S. J. F. Macdonald, J. A. Roper, R. G. Jenkins, R. J. D. Hatley, Emerging therapeutic opportunities for integrin inhibitors. Nat Rev Drug Discov 21, 60–78 (2022).

42. M. H. G. Fonseca, G. P. Furtado, M. R. L. Bezerra, L. Q. Pontes, C. F. C. Fernandes, Boosting half-life and effector functions of therapeutic antibodies by Fc-engineering: An interaction-function review. International Journal of Biological Macromolecules 119, 306–311 (2018).

43. O. Stüve, J. L. Bennett, Pharmacological properties, toxicology and scientific rationale for the use of natalizumab (Tysabri) in inflammatory diseases. CNS Drug Rev 13, 79–95 (2007).

44. T. Rispens, et al., Measurement of serum levels of natalizumab, an immunoglobulin G4 therapeutic monoclonal antibody. Anal Biochem 411, 271– 276 (2011).

45. L. Börnsen, et al., Effect of Natalizumab on Circulating CD4+ T-Cells in Multiple Sclerosis. PLoS One 7, e47578 (2012).

46. B. R. Nielsen, et al., Characterization of naïve, memory and effector T cells in progressive multiple sclerosis. Journal of Neuroimmunology 310, 17–25 (2017).

47. R. Alon, S. W. Feigelson, Chemokine-triggered leukocyte arrest: force-regulated bi-directional integrin activation in quantal adhesive contacts. Curr Opin Cell Biol 24, 670–676 (2012).

